# Common default mode network dysfunction across psychopathologies: A neuroimaging meta-analysis of the n-back working memory paradigm

**DOI:** 10.1101/2020.01.30.927210

**Authors:** Michael C Farruggia, Angela R Laird, Aaron T Mattfeld

**Affiliations:** Interdepartmental Neuroscience Program, Yale University, 333 Cedar Street, New Haven, CT, U.S.; Department of Psychiatry, Yale University School of Medicine, 300 George Street, New Haven, CT 06511, USA; Modern Diet and Physiology Research Center, New Haven, CT 06519, USA; Department of Physics, Florida International University, 11200 SW 8th Street, DM 256, Miami, Fl, U.S.; Department of Psychology, Florida International University, 11200 SW 8th Street, DM 256, Miami, Fl, U.S.; Center for Children and Families, Florida International University, 11200 SW 8th Street, DM 256, Miami, Fl, U.S.

**Keywords:** n-back task, meta-analysis, working memory, psychopathology, ADHD, addiction, depression, bipolar disorder, schizophrenia

## Abstract

The National Institute of Mental Health’s (NIMH) Research Domain Criteria (RDoC) classifies disorders based on shared aspects of behavioral and neurobiological dysfunction. One common behavioral deficit observed in various psychopathologies, namely ADHD, addiction, bipolar disorder, depression, and schizophrenia, is a deficit in working memory performance. However, it is not known to what extent, if any, these disorders share common neurobiological abnormalities that contribute to decrements in performance. The goal of the present study was to examine convergence and divergence of working memory networks across psychopathologies. We used the Activation Likelihood Estimate (ALE) meta-analytic technique to collapse prior data obtained from published studies using the n-back working memory paradigm in individuals with a DSM-criteria diagnosis of the aforementioned disorders. These studies examined areas in the brain that showed increases in activity as a function of working memory-related load compared to a baseline condition, both within subjects and between healthy individuals and those with psychiatric disorder. A meta-analysis of 281 foci covering 81 experiments and 2,629 participants found significant convergence of hyperactivity in medial prefrontal cortex (mPFC) for DSM-diagnosed individuals compared to healthy controls. Foci from ADHD, addiction, bipolar disorder, schizophrenia, and major depression studies contributed to the formation of this cluster. These results provide evidence that default-mode intrusion may constitute a shared seed of dysregulation across multiple psychopathologies, ultimately resulting in poorer working memory performance.

## INTRODUCTION

Deficits in working memory performance are a shared feature across many psychopathologies: depression (1), attention-deficit hyperactivity disorder (ADHD) (2), schizophrenia (3, 4), addiction (5), and bipolar disorder (6). The extent to which the observed impairments are the result of similar neurobiological abnormalities has not been systematically explored. Understanding the shared as well as the unique neurobiological mechanisms that are related to poor working memory performance in different psychopathologies may impact understanding of their pathophysiology, as well as inform the diagnosis and treatment of these diseases. Moreover, this approach seeks to bridge the gap between clinically-derived classification schemes and cognitive neuroscience research outlined by the National Institute of Mental Health’s (NIMH) Research Domain Criteria (RDoC) initiative (7). To our knowledge, no attempts have been made to reconcile the varied functional neuroimaging data that have emerged from research examining working memory across psychopathologies. Patients with schizophrenia, for example, have been shown to exhibit both hypoactivity and hyperactivity in left middle frontal gyrus in response to increased working memory load on the n-back task (3, 8). Similarly, in Major Depressive Disorder (MDD), studies have reported decreases in insula activity during n-back performance (9) while others have reported the opposite effect (10).

To make sense of these disparate results, meta-analytic techniques such as the Activation Likelihood Estimation (ALE) algorithm can be implemented (11, 12). The ALE algorithm provides a statistically rigorous approach to aggregating neuroimaging data that allows researchers to draw inferences using whole-brain peak coordinates (13, 14). The ALE method, aggregating over studies, can be used to determine the likelihood a region contributes to a given contrast of interest (12). This is particularly valuable due to the drawbacks of individual neuroimaging studies, namely low statistical power, high false positive rates, and potential for software errors (15).

The goal of the present study was to examine the convergence and/or divergence of functional neuroimaging findings as it relates to working memory-related load across psychopathologies, namely, ADHD, schizophrenia, addiction, depression, and bipolar disorder. We chose to examine working memory-related load across various permutations of the n-back task, which has been extensively validated as a probe for our psychological construct of interest (16). First, we aggregated all peer-reviewed publications meeting our search criteria. All papers had to include contrasts with patients having a DSM diagnosis of interest. Exceptions were made for the ‘Addiction’ contrast to include illicit substance use more broadly, due to the lack of a pervasive application of DSM criteria in this case. Imaging analyses furthermore had to include a *minimum* n-back contrast of 2-back > rest. That is, a 2-back (or 3-back) > rest contrast was acceptable for our purposes, but not 1-back > rest, 1-back > 0-back, or 3-back with no baseline. Second, we extracted coordinates from all articles reporting within-group contrasts as well as between-group contrasts (e.g., controls > bipolar disorder). Third, we ran each contrast in the GingerALE software to create thresholded ALE images. We examined convergence within psychopathologies to characterize the working memory-related activation patterns specific to each disease. We then examined convergence between psychopathologies to identify regions that showed either shared or unique contributions to the neurobiological differences related to working memory performance.

Differences between psychopathology and control group activation maps may constitute regions that warrant further investigation. Any focus of hyperactivation or hypoactivation across psychopathologies compared to controls may represent functional biomarkers of pathology. In particular, these regions (or region) may play a pivotal role in the pathophysiology of working-memory related deficits commonly observed in ADHD, addiction, bipolar disorder, depression and schizophrenia. Such a finding may suggest a novel target for treatment, a potential new way to predict the onset of mental illness, or a functional consequence resulting from the development of psychiatric illness.

## METHODS

### Criteria Selection for Data Used for Meta-Analysis

We conducted literature searches in both Google Scholar and PubMed utilizing the following terms alone and in combination: ‘n-back,’ ‘working memory,’ ‘2-back,’ and ‘3-back’. We constrained our results by keywords related to our psychopathologies of interest: ‘addiction,’ ‘nicotine,’ ‘cocaine,’ ‘marijuana,’ ‘ecstasy,’ ‘MDMA,’ ‘schizophrenia,’ ‘schizoaffective,’ ‘MDD,’ ‘depression,’ ‘bipolar,’ ‘mania,’ and ‘ADHD.’ Peer reviewed articles were also obtained using BrainMap’s Sleuth 3.0.3 software (17) employing the search procedure: Experiments → Paradigm Class → n-back and Subjects → Diagnosis → Alcoholism, Attention Deficit/ Hyperactivity Disorder, Bipolar Disorder, Depression, Major Depressive Disorder, Schizophrenia, and Substance Use Disorder. The obtained papers were further searched for citations of interest and authors were contacted directly if peak coordinates were missing from their reported analyses.

Studies were grouped into separate diagnostic categories determined by DSM-criteria at the time of their publication. This procedure resulted in the following five categories: Addiction, ADHD, Bipolar Disorder, MDD, and Schizophrenia. We used the DSM-V diagnostic criteria for classification purposes if a disorder was ambiguous. For example, Schizoaffective disorder was classified within the “Schizophrenia” category, as it fits within the DSM-V’s diagnostic category of “Schizophrenia Spectrum and Other Psychotic Disorders.” We included subjects with illicit substance use in the ‘Addiction’ contrast even if they did not have a DSM criteria diagnosis.

We only included experiments using variants (e.g., phonological, visuospatial, emotional) of the n-back paradigm (16) in our meta-analysis. Acceptable experimental conditions included 3-back and 2-back paradigms, while 1-back, 0-back, fixation, and resting conditions were considered acceptable baselines. Preferred baseline conditions were 1-back and 0-back conditions, as subtraction of these conditions allows for removal of common sensory-motor effects associated with subjects’ responses via button press (18). In rare cases, however, fixation or rest conditions were included to mitigate the low power inherent to several between-groups contrasts (i.e., Addiction vs. Healthy Controls). We chose to limit our literature review to experiments that only used the n-back paradigm to hold the task and cognitive process of interest (e.g., working memory) constant while assessing neurobiological variability related to different psychopathologies.

Neuroimaging experiments were included only if brain scans were acquired using whole-brain functional magnetic resonance imaging (fMRI) or positron emission tomography (PET). Only studies that reported whole-brain analyses, as opposed to region of interest (ROI) analyses, with coordinates listed in standard stereotactic space (MNI or Talairach/Tournoux), were included in our subsequent ALE analyses. All MNI coordinates were converted to Talairach coordinates (19) using a transformation created by Lacadie et al. (20).

Our minimum statistical criteria for inclusion consisted of studies with a significance threshold of *p* < 0.001 uncorrected in at least 8 individuals (Eickhoff, personal communication, 2016). In total, we identified 160 experiments that matched our criteria, comprised of 54 control, 20 Addiction, 17 ADHD, 25 Bipolar, 15 MDD, and 29 schizophrenia experiments. These experiments reported 1,603 brain activation foci obtained from a total of 4,509 participants. Our search procedure was concluded in October of 2019.

### Activation Likelihood Estimation Algorithm

To examine the brain regions activated during the n-back task across addiction, ADHD, bipolar disorder, MDD, and schizophrenia psychopathologies, we performed a coordinate-based meta-analysis using GingerALE v3.0.2 (17). GingerALE’s revised ALE algorithm creates a statistical map using the supplied peak coordinates to estimate the likelihood of activation of each voxel in the brain. Activation foci are viewed as centers of 3-D Gaussian probability distribution functions, which are used to estimate the probability that at least one of the activation foci in the dataset actually lies within a given voxel (12) – these probabilities are known as Activation Likelihood Estimate (ALE) values. Importantly, the ALE algorithm weighs the between-subject variance by the number of subjects in each study, such that larger studies are associated with narrower Gaussian distributions than smaller studies. Maps were then created using the voxel-wise ALE values for each contrast. The resulting ALE maps were thresholded at *p* < 0.01 uncorrected and then subjected to a permutation test (1000 replications) with a cluster threshold value *p* < 0.01 FWE (family-wise error).

### Contrasts

We conducted multiple ALE meta-analyses, both within and between-groups. Initially, we characterized the n-back working memory network separately for healthy controls, all psychopathologies collapsed, and for each individual psychopathology (i.e. within-group contrasts). In the healthy controls contrast, we examined coordinates from 54 experiments, comprising a total of 1,040 healthy control participants and 561 foci. We only included publications that obtained data from healthy participants and were additionally used in the within-group ‘psychopathologies’ contrasts. In the psychopathologies contrasts, we included data from a total of 106 experiments: 20 Addiction, 17 ADHD, 25 Bipolar, 15 MDD, and 29 schizophrenia. These experiments included a total of 1,042 foci and 3,469 participants. Next, we evaluated brain networks that were hyperactive in controls compared to all combined psychopathologies during the n-back task (i.e. between-group contrasts) and also compared individually as follow-up. A total of 73 experiments fit eligibility criteria for this analysis: 8 Controls > Addiction, 12 Controls > ADHD, 15 Controls > Bipolar, 11 Controls > MDD experiments, and 27 Controls > Schizophrenia; resulting in 336 foci from 2,788 participants. Lastly, we examined the brain networks that were hyperactive during the n-back task across all psychopathologies combined and individually, in comparison to healthy controls. For this analysis, we isolated 81 experiments that matched our criteria: 32 schizophrenia > controls, 18 bipolar > controls, 11 MDD > controls, 11 addiction > controls, and 9 ADHD > controls; consisting of 281 foci obtained from 2,629 participants. A full list of studies included in our within- and between-groups contrasts are available in Tables 1 and 2, respectively. For all analyses and contrasts, we report anatomical labels (Talairach Nearest Grey Matter) of the weighted center (x,y,z) of each obtained cluster. Clusters were overlaid onto the standard “Colin” brain in Talairach space (21) using Mango v. 4.1 software (http://ric.uthscsa.edu/mango/).

**TABLE 1.**
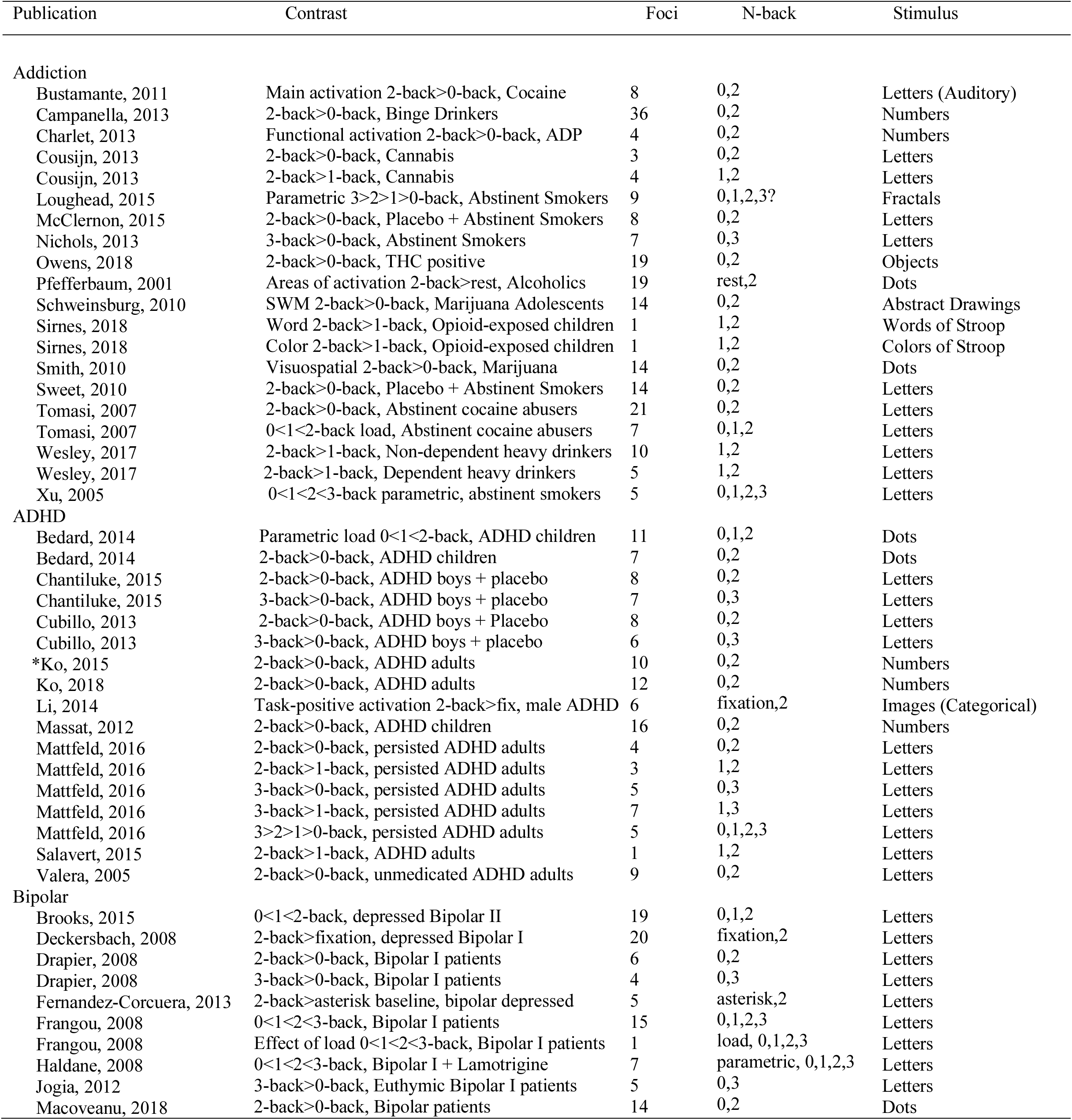
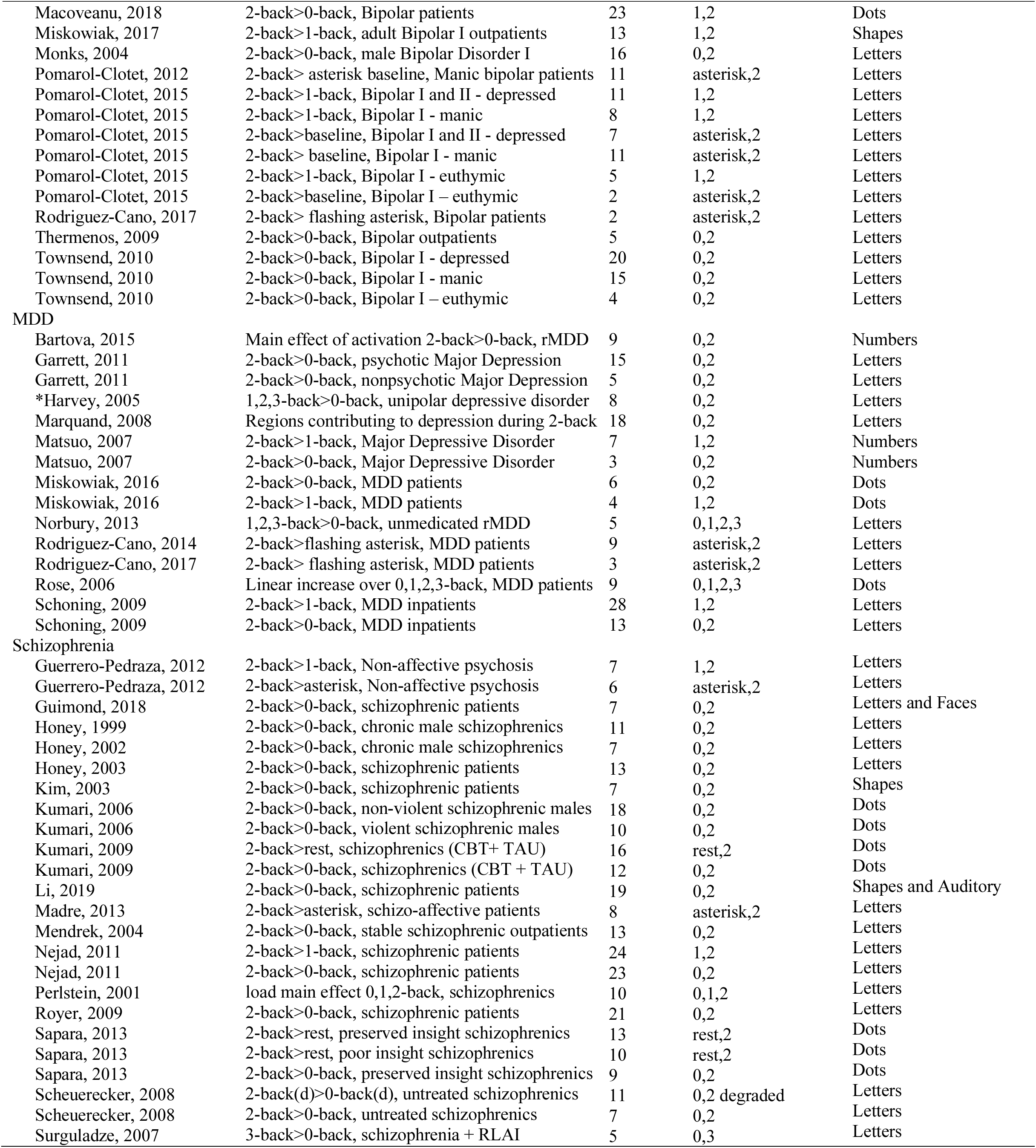
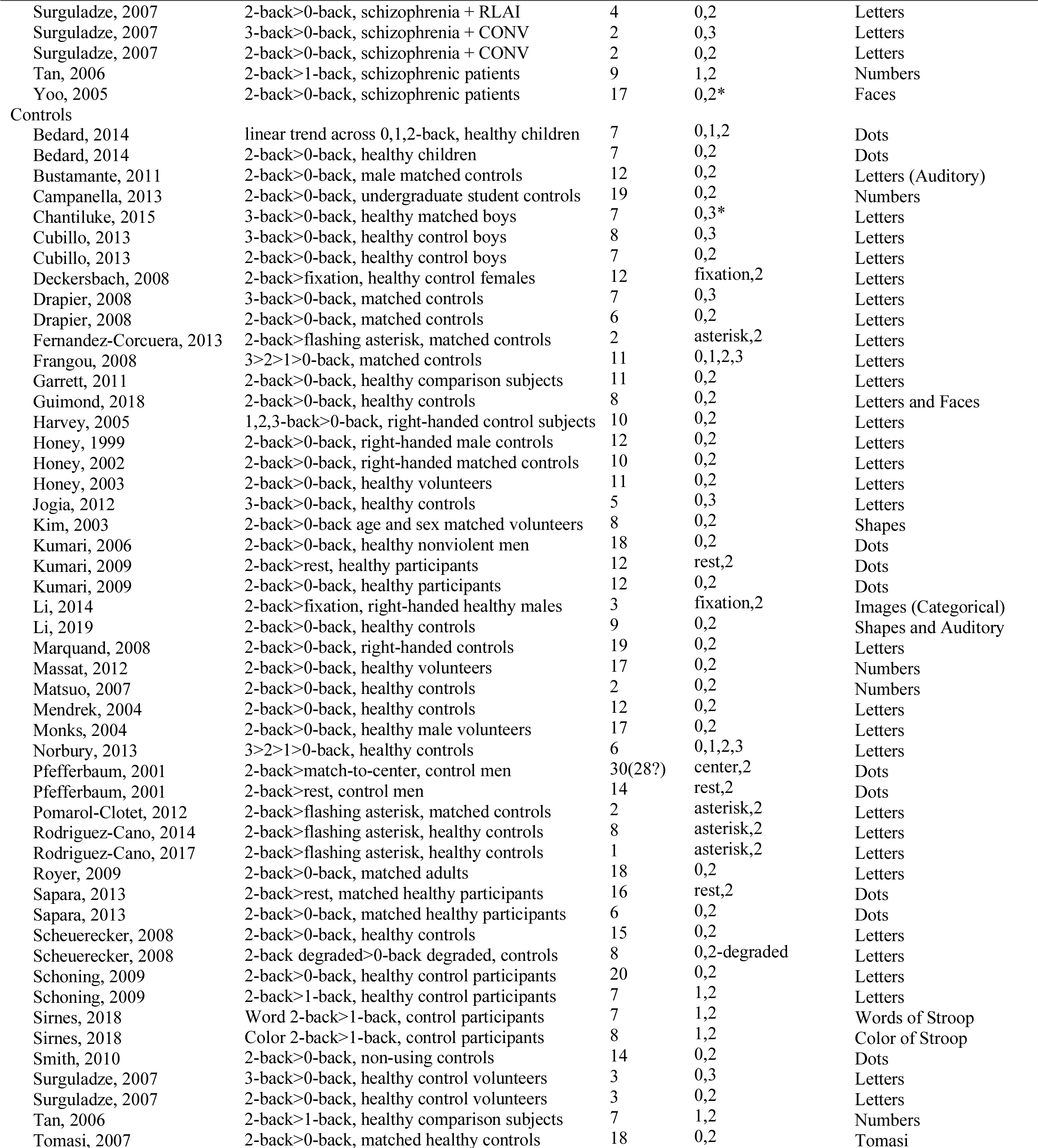
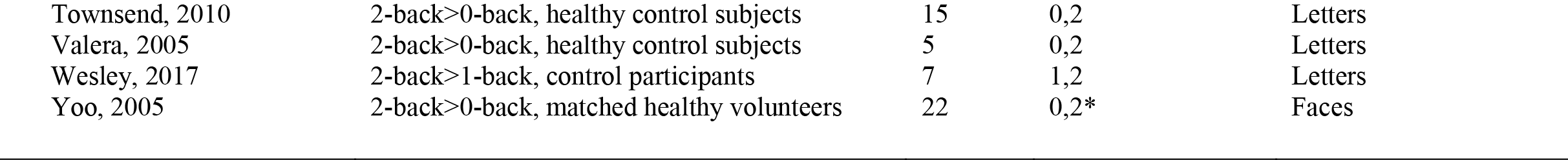
WITHIN-GROUPS CONTRASTS

| Publication | Contrast | Foci | N-back | Stimulus |
| --- | --- | --- | --- | --- |
| **Addiction** |  |  |  |  |
| Bustamante, 2011 | Main activation 2-back>0-back, Cocaine | 8 | 0,2 | Letters (Auditory) |
| Campanella, 2013 | 2-back>0-back, Binge Drinkers | 36 | 0,2 | Numbers |
| Charlet, 2013 | Functional activation 2-back>0-back, ADP | 4 | 0,2 | Numbers |
| Cousijn, 2013 | 2-back>0-back, Cannabis | 3 | 0,2 | Letters |
| Cousijn, 2013 | 2-back>1-back, Cannabis | 4 | 1,2 | Letters |
| Loughead, 2015 | Parametric 3>2>1>0-back, Abstinent Smokers | 9 | 0,1,2,3? | Fractals |
| McClernon, 2015 | 2-back>0-back, Placebo + Abstinent Smokers | 8 | 0,2 | Letters |
| Nichols, 2013 | 3-back>0-back, Abstinent Smokers | 7 | 0,3 | Letters |
| Owens, 2018 | 2-back>0-back, THC positive | 19 | 0,2 | Objects |
| Pfefferbaum, 2001 | Areas of activation 2-back>rest, Alcoholics | 19 | rest,2 | Dots |
| Schweinsburg, 2010 | SWM 2-back>0-back, Marijuana Adolescents | 14 | 0,2 | Abstract Drawings |
| Sirnes, 2018 | Word 2-back>1-back, Opioid-exposed children | 1 | 1,2 | Words of Stroop |
| Sirnes, 2018 | Color 2-back>1-back, Opioid-exposed children | 1 | 1,2 | Colors of Stroop |
| Smith, 2010 | Visuospatial 2-back>0-back, Marijuana | 14 | 0,2 | Dots |
| Sweet, 2010 | 2-back>0-back, Placebo + Abstinent Smokers | 14 | 0,2 | Letters |
| Tomasi, 2007 | 2-back>0-back, Abstinent cocaine abusers | 21 | 0,2 | Letters |
| Tomasi, 2007 | 0<1<2-back load, Abstinent cocaine abusers | 7 | 0,1,2 | Letters |
| Wesley, 2017 | 2-back>1-back, Non-dependent heavy drinkers | 10 | 1,2 | Letters |
| Wesley, 2017 | 2-back>1-back, Dependent heavy drinkers | 5 | 1,2 | Letters |
| Xu, 2005 | 0<1<2<3-back parametric, abstinent smokers | 5 | 0,1,2,3 | Letters |
| **ADHD** |  |  |  |  |
| Bedard, 2014 | Parametric load 0<1<2-back, ADHD children | 11 | 0,1,2 | Dots |
| Bedard, 2014 | 2-back>0-back, ADHD children | 7 | 0,2 | Dots |
| Chantiluke, 2015 | 2-back>0-back, ADHD boys + placebo | 8 | 0,2 | Letters |
| Chantiluke, 2015 | 3-back>0-back, ADHD boys + placebo | 7 | 0,3 | Letters |
| Cubillo, 2013 | 2-back>0-back, ADHD boys + Placebo | 8 | 0,2 | Letters |
| Cubillo, 2013 | 3-back>0-back, ADHD boys + placebo | 6 | 0,3 | Letters |
| *Ko, 2015 | 2-back>0-back, ADHD adults | 10 | 0,2 | Numbers |
| Ko, 2018 | 2-back>0-back, ADHD adults | 12 | 0,2 | Numbers |
| Li, 2014 | Task-positive activation 2-back>fix, male ADHD | 6 | fixation,2 | Images (Categorical) |
| Massat, 2012 | 2-back>0-back, ADHD children | 16 | 0,2 | Numbers |
| Mattfeld, 2016 | 2-back>0-back, persisted ADHD adults | 4 | 0,2 | Letters |
| Mattfeld, 2016 | 2-back>1-back, persisted ADHD adults | 3 | 1,2 | Letters |
| Mattfeld, 2016 | 3-back>0-back, persisted ADHD adults | 5 | 0,3 | Letters |
| Mattfeld, 2016 | 3-back>1-back, persisted ADHD adults | 7 | 1,3 | Letters |
| Mattfeld, 2016 | 3>2>1>0-back, persisted ADHD adults | 5 | 0,1,2,3 | Letters |
| Salavert, 2015 | 2-back>1-back, ADHD adults | 1 | 1,2 | Letters |
| Valera, 2005 | 2-back>0-back, unmedicated ADHD adults | 9 | 0,2 | Letters |
| **Bipolar** |  |  |  |  |
| Brooks, 2015 | 0<1<2-back, depressed Bipolar II | 19 | 0,1,2 | Letters |
| Deckersbach, 2008 | 2-back>fixation, depressed Bipolar I | 20 | fixation,2 | Letters |
| Drapier, 2008 | 2-back>0-back, Bipolar I patients | 6 | 0,2 | Letters |
| Drapier, 2008 | 3-back>0-back, Bipolar I patients | 4 | 0,3 | Letters |
| Fernandez-Corcuera, 2013 | 2-back>asterisk baseline, bipolar depressed | 5 | asterisk,2 | Letters |
| Frangou, 2008 | 0<1<2<3-back, Bipolar I patients | 15 | 0,1,2,3 | Letters |
| Frangou, 2008 | Effect of load 0<1<2<3-back, Bipolar I patients | 1 | load, 0,1,2,3 | Letters |
| Haldane, 2008 | 0<1<2<3-back, Bipolar I + Lamotrigine | 7 | parametric, 0,1,2,3 | Letters |
| Jogia, 2012 | 3-back>0-back, Euthymic Bipolar I patients | 5 | 0,3 | Letters |
| Macoveanu, 2018 | 2-back>0-back, Bipolar patients | 14 | 0,2 | Dots |
| Macoveanu, 2018 | 2-back>0-back, Bipolar patients | 23 | 1,2 | Dots |
| Miskowiak, 2017 | 2-back>1-back, adult Bipolar I outpatients | 13 | 1,2 | Shapes |
| Monks, 2004 | 2-back>0-back, male Bipolar Disorder I | 16 | 0,2 | Letters |
| Pomarol-Clotet, 2012 | 2-back> asterisk baseline, Manic bipolar patients | 11 | asterisk,2 | Letters |
| Pomarol-Clotet, 2015 | 2-back>1-back, Bipolar I and II - depressed | 11 | 1,2 | Letters |
| Pomarol-Clotet, 2015 | 2-back>1-back, Bipolar I - manic | 8 | 1,2 | Letters |
| Pomarol-Clotet, 2015 | 2-back>baseline, Bipolar I and II - depressed | 7 | asterisk,2 | Letters |
| Pomarol-Clotet, 2015 | 2-back> baseline, Bipolar I - manic | 11 | asterisk,2 | Letters |
| Pomarol-Clotet, 2015 | 2-back>1-back, Bipolar I - euthymic | 5 | 1,2 | Letters |
| Pomarol-Clotet, 2015 | 2-back>baseline, Bipolar I – euthymic | 2 | asterisk,2 | Letters |
| Rodriguez-Cano, 2017 | 2-back> flashing asterisk, Bipolar patients | 2 | asterisk,2 | Letters |
| Thermenos, 2009 | 2-back>0-back, Bipolar outpatients | 5 | 0,2 | Letters |
| Townsend, 2010 | 2-back>0-back, Bipolar I - depressed | 20 | 0,2 | Letters |
| Townsend, 2010 | 2-back>0-back, Bipolar I - manic | 15 | 0,2 | Letters |
| Townsend, 2010 | 2-back>0-back, Bipolar I – euthymic | 4 | 0,2 | Letters |
| MDD |  |  |  |  |
| Bartova, 2015 | Main effect of activation 2-back>0-back, rMDD | 9 | 0,2 | Numbers |
| Garrett, 2011 | 2-back>0-back, psychotic Major Depression | 15 | 0,2 | Letters |
| Garrett, 2011 | 2-back>0-back, nonpsychotic Major Depression | 5 | 0,2 | Letters |
| *Harvey, 2005 | 1,2,3-back>0-back, unipolar depressive disorder | 8 | 0,2 | Letters |
| Marquand, 2008 | Regions contributing to depression during 2-back | 18 | 0,2 | Letters |
| Matsuo, 2007 | 2-back>1-back, Major Depressive Disorder | 7 | 1,2 | Numbers |
| Matsuo, 2007 | 2-back>0-back, Major Depressive Disorder | 3 | 0,2 | Numbers |
| Miskowiak, 2016 | 2-back>0-back, MDD patients | 6 | 0,2 | Dots |
| Miskowiak, 2016 | 2-back>1-back, MDD patients | 4 | 1,2 | Dots |
| Norbury, 2013 | 1,2,3-back>0-back, unmedicated rMDD | 5 | 0,1,2,3 | Letters |
| Rodriguez-Cano, 2014 | 2-back>flashing asterisk, MDD patients | 9 | asterisk,2 | Letters |
| Rodriguez-Cano, 2017 | 2-back> flashing asterisk, MDD patients | 3 | asterisk,2 | Letters |
| Rose, 2006 | Linear increase over 0,1,2,3-back, MDD patients | 9 | 0,1,2,3 | Dots |
| Schoning, 2009 | 2-back>1-back, MDD inpatients | 28 | 1,2 | Letters |
| Schoning, 2009 | 2-back>0-back, MDD inpatients | 13 | 0,2 | Letters |
| Schizophrenia |  |  |  |  |
| Guerrero-Pedraza, 2012 | 2-back>1-back, Non-affective psychosis | 7 | 1,2 | Letters |
| Guerrero-Pedraza, 2012 | 2-back>asterisk, Non-affective psychosis | 6 | asterisk,2 | Letters |
| Guimond, 2018 | 2-back>0-back, schizophrenic patients | 7 | 0,2 | Letters and Faces |
| Honey, 1999 | 2-back>0-back, chronic male schizophrenics | 11 | 0,2 | Letters |
| Honey, 2002 | 2-back>0-back, chronic male schizophrenics | 7 | 0,2 | Letters |
| Honey, 2003 | 2-back>0-back, schizophrenic patients | 13 | 0,2 | Letters |
| Kim, 2003 | 2-back>0-back, schizophrenic patients | 7 | 0,2 | Shapes |
| Kumari, 2006 | 2-back>0-back, non-violent schizophrenic males | 18 | 0,2 | Dots |
| Kumari, 2006 | 2-back>0-back, violent schizophrenic males | 10 | 0,2 | Dots |
| Kumari, 2009 | 2-back>rest, schizophrenics (CBT+ TAU) | 16 | rest,2 | Dots |
| Kumari, 2009 | 2-back>0-back, schizophrenics (CBT + TAU) | 12 | 0,2 | Dots |
| Li, 2019 | 2-back>0-back, schizophrenic patients | 19 | 0,2 | Shapes and Auditory |
| Madre, 2013 | 2-back>asterisk, schizo-affective patients | 8 | asterisk,2 | Letters |
| Mendrek, 2004 | 2-back>0-back, stable schizophrenic outpatients | 13 | 0,2 | Letters |
| Nejad, 2011 | 2-back>1-back, schizophrenic patients | 24 | 1,2 | Letters |
| Nejad, 2011 | 2-back>0-back, schizophrenic patients | 23 | 0,2 | Letters |
| Perlstein, 2001 | load main effect 0,1,2-back, schizophrenics | 10 | 0,1,2 | Letters |
| Royer, 2009 | 2-back>0-back, schizophrenic patients | 21 | 0,2 | Letters |
| Sapara, 2013 | 2-back>rest, preserved insight schizophrenics | 13 | rest,2 | Dots |
| Sapara, 2013 | 2-back>rest, poor insight schizophrenics | 10 | rest,2 | Dots |
| Sapara, 2013 | 2-back>0-back, preserved insight schizophrenics | 9 | 0,2 | Dots |
| Scheuerecker, 2008 | 2-back(d)>0-back(d), untreated schizophrenics | 11 | 0,2 degraded | Letters |
| Scheuerecker, 2008 | 2-back>0-back, untreated schizophrenics | 7 | 0,2 | Letters |
| Surguladze, 2007 | 3-back>0-back, schizophrenia + RLAI | 5 | 0,3 | Letters |
| Surguladze, 2007 | 2-back>0-back, schizophrenia + RLAI | 4 | 0,2 | Letters |
| Surguladze, 2007 | 3-back>0-back, schizophrenia + CONV | 2 | 0,3 | Letters |
| Surguladze, 2007 | 2-back>0-back, schizophrenia + CONV | 2 | 0,2 | Letters |
| Tan, 2006 | 2-back>1-back, schizophrenic patients | 9 | 1,2 | Numbers |
| Yoo, 2005 | 2-back>0-back, schizophrenic patients | 17 | 0,2* | Faces |
| Controls |  |  |  |  |
| Bedard, 2014 | linear trend across 0,1,2-back, healthy children | 7 | 0,1,2 | Dots |
| Bedard, 2014 | 2-back>0-back, healthy children | 7 | 0,2 | Dots |
| Bustamante, 2011 | 2-back>0-back, male matched controls | 12 | 0,2 | Letters (Auditory) |
| Campanella, 2013 | 2-back>0-back, undergraduate student controls | 19 | 0,2 | Numbers |
| Chantiluke, 2015 | 3-back>0-back, healthy matched boys | 7 | 0,3* | Letters |
| Cubillo, 2013 | 3-back>0-back, healthy control boys | 8 | 0,3 | Letters |
| Cubillo, 2013 | 2-back>0-back, healthy control boys | 7 | 0,2 | Letters |
| Deckersbach, 2008 | 2-back>fixation, healthy control females | 12 | fixation,2 | Letters |
| Drapier, 2008 | 3-back>0-back, matched controls | 7 | 0,3 | Letters |
| Drapier, 2008 | 2-back>0-back, matched controls | 6 | 0,2 | Letters |
| Fernandez-Corcuera, 2013 | 2-back>flashing asterisk, matched controls | 2 | asterisk,2 | Letters |
| Frangou, 2008 | 3>2>1>0-back, matched controls | 11 | 0,1,2,3 | Letters |
| Garrett, 2011 | 2-back>0-back, healthy comparison subjects | 11 | 0,2 | Letters |
| Guimond, 2018 | 2-back>0-back, healthy controls | 8 | 0,2 | Letters and Faces |
| Harvey, 2005 | 1,2,3-back>0-back, right-handed control subjects | 10 | 0,2 | Letters |
| Honey, 1999 | 2-back>0-back, right-handed male controls | 12 | 0,2 | Letters |
| Honey, 2002 | 2-back>0-back, right-handed matched controls | 10 | 0,2 | Letters |
| Honey, 2003 | 2-back>0-back, healthy volunteers | 11 | 0,2 | Letters |
| Jogia, 2012 | 3-back>0-back, healthy controls | 5 | 0,3 | Letters |
| Kim, 2003 | 2-back>0-back age and sex matched volunteers | 8 | 0,2 | Shapes |
| Kumari, 2006 | 2-back>0-back, healthy nonviolent men | 18 | 0,2 | Dots |
| Kumari, 2009 | 2-back>rest, healthy participants | 12 | rest,2 | Dots |
| Kumari, 2009 | 2-back>0-back, healthy participants | 12 | 0,2 | Dots |
| Li, 2014 | 2-back>fixation, right-handed healthy males | 3 | fixation,2 | Images (Categorical) |
| Li, 2019 | 2-back>0-back, healthy controls | 9 | 0,2 | Shapes and Auditory |
| Marquand, 2008 | 2-back>0-back, right-handed controls | 19 | 0,2 | Letters |
| Massat, 2012 | 2-back>0-back, healthy volunteers | 17 | 0,2 | Numbers |
| Matsuo, 2007 | 2-back>0-back, healthy controls | 2 | 0,2 | Numbers |
| Mendrek, 2004 | 2-back>0-back, healthy controls | 12 | 0,2 | Letters |
| Monks, 2004 | 2-back>0-back, healthy male volunteers | 17 | 0,2 | Letters |
| Norbury, 2013 | 3>2>1>0-back, healthy controls | 6 | 0,1,2,3 | Letters |
| Pfefferbaum, 2001 | 2-back>match-to-center, control men | 30(28?) | center,2 | Dots |
| Pfefferbaum, 2001 | 2-back>rest, control men | 14 | rest,2 | Dots |
| Pomarol-Clotet, 2012 | 2-back>flashing asterisk, matched controls | 2 | asterisk,2 | Letters |
| Rodriguez-Cano, 2014 | 2-back>flashing asterisk, healthy controls | 8 | asterisk,2 | Letters |
| Rodriguez-Cano, 2017 | 2-back>flashing asterisk, healthy controls | 1 | asterisk,2 | Letters |
| Royer, 2009 | 2-back>0-back, matched adults | 18 | 0,2 | Letters |
| Sapara, 2013 | 2-back>rest, matched healthy participants | 16 | rest,2 | Dots |
| Sapara, 2013 | 2-back>0-back, matched healthy participants | 6 | 0,2 | Dots |
| Scheuerecker, 2008 | 2-back>0-back, healthy controls | 15 | 0,2 | Letters |
| Scheuerecker, 2008 | 2-back degraded>0-back degraded, controls | 8 | 0,2-degraded | Letters |
| Schoning, 2009 | 2-back>0-back, healthy control participants | 20 | 0,2 | Letters |
| Schoning, 2009 | 2-back>1-back, healthy control participants | 7 | 1,2 | Letters |
| Sirnes, 2018 | Word 2-back>1-back, control participants | 7 | 1,2 | Words of Stroop |
| Sirnes, 2018 | Color 2-back>1-back, control participants | 8 | 1,2 | Color of Stroop |
| Smith, 2010 | 2-back>0-back, non-using controls | 14 | 0,2 | Dots |
| Surguladze, 2007 | 3-back>0-back, healthy control volunteers | 3 | 0,3 | Letters |
| Surguladze, 2007 | 2-back>0-back, healthy control volunteers | 3 | 0,2 | Letters |
| Tan, 2006 | 2-back>1-back, healthy comparison subjects | 7 | 1,2 | Numbers |
| Tomasi, 2007 | 2-back>0-back, matched healthy controls | 18 | 0,2 | Tomasi |
| Townsend, 2010 | 2-back>0-back, healthy control subjects | 15 | 0,2 | Letters |
| Valera, 2005 | 2-back>0-back, healthy control subjects | 5 | 0,2 | Letters |
| Wesley, 2017 | 2-back>1-back, control participants | 7 | 1,2 | Letters |
| Yoo, 2005 | 2-back>0-back, matched healthy volunteers | 22 | 0,2* | Faces |

**TABLE 2.** BETWEEN-GROUPS CONTRASTS

| Publication | Contrast | Foci | N-back | Stimulus |
| --- | --- | --- | --- | --- |
| **Addiction > Controls** |  |  |  |  |
| Campanella, 2013 | 2-back>0-back, binge drinkers>controls | 2 | 0,2 | Numbers |
| Charlet, 2013 | 2-back>0-back, detoxified ADP>controls | 12 | 0,2 | Numbers |
| Padula, 2007 | 2-back>0-back, abstinent MJ>controls | 3 | 0,2 | Abstract Shapes |
| Sirnes, 2018 | 2-back>1-back, opioid-exposed children | 2 | 1,2 | Colors |
| Sirnes, 2018 | 2-back>1-back, opioid-exposed children | 1 | 1,2 | Words |
| Smith, 2010 | 2-back>0-back, MJ>controls | 3 | 0,2 | Dots |
| Tomasi, 2007 | 2-back>0-back, Cocaine>controls | 11 | 0,2 | Letters |
| Tomasi, 2007 | 2>1>0-back, Cocaine>controls | 1 | 0,1,2 load | Letters |
| Wesley, 2017 | 2-back>1-back, Alcohol-dependent>Controls | 1 | 1,2 | Letters |
| **Addiction < Controls** |  |  |  |  |
| Bustamante, 2011 | 2-back>0-back, controls>cocaine-dependent | 1 | 0,2 | Letters (Auditory) |
| Daumann, 2003 | 2-back>rest, controls>heavy MDMA users | 1 | rest,2 | Letters |
| Daumann(2), 2003 | 2-back>rest, controls> heavy MDMA users | 1 | rest,2 | Letters |
| Livny, 2018 | 2-back>0-back, controls>synthetic cannabinoid | 1 | 0,2 | Letters |
| Livny, 2018 | 2-back>1-back, controls> synthetic cannabinoid | 2 | 1,2 | Letters |
| Pfefferbaum, 2001 | 2-back>rest, controls>chronic alcoholics | 24 | rest,2 | Dots |
| Tomasi, 2007 | 2-back>0-back, controls>cocaine abusers | 10 | 0,2 | Letters |
| Tomasi, 2007 | 2>1>0-back, controls>cocaine abusers | 2 | 0,1,2 load | Letters |
| **ADHD > Controls** |  |  |  |  |
| Bedard, 2014 | 2-back>0-back, ADHD youth>controls | 2 | 0,2 | Dots |
| Cubillo, 2013 | 3>2>1>0-back, ADHD+MPH boys>controls | 1 | 0,1,2,3 load | Letters |
| Ko, 2015 | 2-back>0-back, ADHD adults>controls | 2 | 0,2 | Numbers |
| Ko, 2018 | 2-back>0-back, ADHD adults>controls | 3 | 0,2 | Numbers |
| Li, 2014 | 2-back>fixation, ADHD males>controls | 3 | fixation,2 | Images (Categorical) |
| Mattfeld, 2016 | 2-back>1-back, ADHD adults>controls | 1 | 1,2 | Letters |
| Mattfeld, 2016 | 3-back>1-back, ADHD adults>controls | 1 | 1,3 | Letters |
| Mattfeld, 2016 | 3-back>2-back, ADHD adults>controls | 2 | 3,2 | Letters |
| Salavert, 2015 | 2-back>asterisk, ADHD adult patients>controls | 1 | asterisk,2 | Letters |
| **ADHD < Controls** |  |  |  |  |
| Bayerl, 2010 | 2-back>0-back, controls>adult ADHD patients | 1 | 0,2 | Letters |
| Brown, 2012 | 2-back>0-back, controls>male ADHD+BPD | 2 | 0,2 | Letters |
| Cubillo, 2013 | 3>2>1>0-back, controls> ADHD+MPH boys | 1 | 0,1,2,3 load | Letters |
| Cubillo, 2013 | 3>2>1>0-back, controls>ADHD+ATX boys | 1 | 0,1,2,3 load | Letters |
| Kobel, 2009 | 3+2>0-back, controls>ADHD boys | 5 | 2,3* | Numbers |
| Mattfeld, 2016 | 2-back>0-back, controls>ADHD adults | 2 | 0,2 | Letters |
| Mattfeld, 2016 | 3-back>0-back, controls>ADHD adults | 1 | 0,3 | Letters |
| Mattfeld, 2016 | 3>2>1>0-back, controls>ADHD adults | 1 | 0,1,2,3 | Letters |
| Mattfeld, 2016 | 3>2>1>0-back, controls>impaired ADHD adults | 5 | 0,1,2,3 | Letters |
| Valera, 2005 | 2-back>0-back, controls>ADHD adults | 2 | 0,2 | Letters |
| Valera, 2005 | 2-back>0-back, controls>ADHD-LD adults | 1 | 0,2 | Letters |
| Valera, 2010 | 2-back>0-back, controls> ADHD adults | 2 | 0,2 | Letters |
| **Bipolar > Controls** |  |  |  |  |
| Adler, 2004 | 2-back>0-back, Bipolar patients>controls | 8 | 0,2 | Numbers |
| Alonso-Lana, 2016 | 2-back>asterisk, euthymic bipolar>controls | 1 | asterisk,2 | Letters |
| Alonso-Lana, 2019 | 2-back>asterisk, manic bipolar>controls | 1 | asterisk,2 | Letters |
| Alonso-Lana, 2019 | 2-back>asterisk, euthymic bipolar>controls | 1 | asterisk,2 | Letters |
| Alonso-Lana, 2019 | 2-back>1-back, manic bipolar>controls | 1 | 1,2 | Letters |
| Deckersbach, 2008 | 2-back>fixation, Bipolar I disorder>control | 4 | fixation,2 | Letters |
| Dima, 2016 | 3-back>0-back, Bipolar patients>control | 3 | 0,3 | Letters |
| Drapier, 2008 | 2-back>0-back, Bipolar I patients>controls | 2 | 0,2 | Letters |
| Fernandez-Corcuera, 2013 | 2-back>asterisk, Bipolar depressed>controls | 1 | asterisk,2 | Letters |
| Frangou, 2017 | 3-back>0-back, BD I patients>relatives+controls | 1 | 0,3 | Letters |
| Jogia, 2012 | 3-back>0-back, euthymic BD I>Controls | 4 | 0,3 | Letters |
| Monks, 2004 | 2-back>0-back, Bipolar disorder I>Controls | 3 | 0,2 | Letters |
| Pomarol-Clotet, 2012 | 2-back>asterisk, manic bipolar>controls | 3 | asterisk,2 | Letters |
| Pomarol-Clotet, 2015 | 2-back>asterisk, Bipolar I manic>controls | 1 | asterisk,2 | Letters |
| Pomarol-Clotet, 2015 | 2-back>asterisk, Bipolar I/II depressed>controls | 1 | asterisk,2 | Letters |
| Pomarol-Clotet, 2015 | 2-back>asterisk, Bipolar I euthymic> controls | 1 | asterisk,2 | Letters |
| Rodriguez-Cano, 2017 | 2-back>asterisk, Bipolar I patients>controls | 1 | asterisk,2 | Letters |
| Thermenos, 2009 | 2-back>0-back, bipolar outpatients>controls | 1 | 0,2 | Letters |
| Bipolar < Controls |  |  |  |  |
| Alonso-Lana, 2019 | 2-back>asterisk, controls> manic bipolar | 2 | asterisk,2 | Letters |
| Alonso-Lana, 2019 | 2-back>1-back, controls>euthymic bipolar | 1 | 1,2 | Letters |
| Brooks, 2015 | 2>1>0-back(load), controls>Bipolar II depressed | 8 | 0,1,2 | Letters |
| Dima, 2016 | 3-back>0-back, controls>Bipolar patients | 4 | 0,3 | Letters |
| Fernandez-Corcuera, 2013 | 2-back>asterisk, controls>Bipolar depressed | 3 | asterisk,2 | Letters |
| Frangou, 2017 | 3-back>0-back, relatives+controls> BD I patients | 1 | 0,3 | Letters |
| Jogia, 2012 | 3-back>0-back, controls>euthymic BD I | 1 | 0,3 | Letters |
| Monks, 2004 | 2-back>0-back, controls>bipolar disorder I | 7 | 0,2 | Letters |
| Moser, 2018 | 2-back>0-back, controls>bipolar disorder | 3 | 0,2 | Objects |
| Meusel, 2013 | 2-back>0-back, controls>MDD+BD | 1 | 0,2 | Abstract Objects |
| Pomarol-Clotet, 2012 | 2-back>asterisk, controls>manic bipolar | 3 | asterisk,2 | Letters |
| Pomarol-Clotet, 2015 | 2-back>asterisk, controls> Bipolar I manic | 6 | asterisk,2 | Letters |
| Pomarol-Clotet, 2015 | 2-back>asterisk, controls>Bipolar I/II depressed | 3 | asterisk,2 | Letters |
| Rodriguez-Cano, 2017 | 2-back>asterisk, controls>Bipolar I patients | 2 | asterisk,2 | Letters |
| Thermenos, 2009 | 2-back>0-back, controls> bipolar outpatients | 1 | 0,2 | Letters |
| MDD > Controls |  |  |  |  |
| Bartova, 2015 | 2-back>0-back, remitted MDD patients>controls | 4 | 0,2 | Numbers |
| Fitzgerald, 2008 | 2-back>0-back, MDD>healthy controls | 20 | 0,2 | Letters |
| Harvey, 2005 | 2-back>0-back, nonpsychotic MDE>controls | 3 | 0,2 | Letters |
| Matsuo, 2007 | 2-back>1-back, recurrent MDD>healthy controls | 2 | 1,2 | Numbers |
| Nord, 2018 | 3-back>1-back, MDD patients>controls | 1 | 1,3 | Letters |
| Nord, 2018 | 3-back>1-back, MDD patients>relatives | 12 | 1,3 | Letters |
| Rodriguez-Cano, 2014 | 2-back>asterisk, MDD>controls | 1 | asterisk,2 | Letters |
| Rodriguez-Cano, 2017 | 2-back>asterisk, MDD>controls | 1 | asterisk,2 | Letters |
| Rose, 2006 | 3>2>1>0-back (linear increase), MDD>Controls | 1 | 0,1,2,3 | Dots |
| Schoning, 2009 | 2-back>0-back, euthymic MDD>Controls | 8 | 0,2 | Letters |
| Yüksel, 2018 | 3-back>2-back, MDD patients>Controls | 1 | 2,3 | Letters |
| MDD < Controls |  |  |  |  |
| Barch, 2003 | 2-back>fixation, controls>unipolar MDD | 4 | fixation,2 | Words and Faces |
| Dumas, 2014 | 2-back>0-back, controls>MDD old adults | 3 | 0,2 | Verbal |
| Frangou, 2017 | 3-back>0-back, controls>MDD | 1 | 0,3 | Letters |
| Ionescu, 2015 | 2-back>1-back, controls>treatment-resistant MD | 3 | 1,2 | Numbers |
| Nord, 2018 | 3-back>1-back, controls>MDD patients | 15 | 1,3 | Letters |
| Nord, 2018 | 3-back>1-back, relatives>MDD patients | 6 | 1,3 | Letters |
| Rodriguez-Cano, 2014 | 2-back>asterisk, controls>MDD | 4 | asterisk,2 | Letters |
| Rodriguez-Cano, 2017 | 2-back>asterisk, controls>MDD | 1 | asterisk,2 | Letters |
| Walsh, 2007 | 3,2,1>0-back, controls>MDD | 2 | 0,1,2,3 | Letters |
| Yüksel, 2018 | 2-back>0-back, controls>MDD patients | 1 | 0,2 | Letters |
| Yüksel, 2018 | 3-back>0-back, controls>MDD patients | 2 | 0,3 | Letters |
| Schizophrenia > Controls |  |  |  |  |
| Bor, 2011 | 2-back>0-back, schizophrenia>controls | 3 | 0,2 | Letters and Dots (s) |
| Bor, 2011 | 2-back>0-back, schizophrenia>controls | 1 | 0,2 | Letters and Dots (v) |
| Becerril, 2010 | 2-back>fixation, schizophrenia>controls | 6 | fixation,2 | Faces |
| Callicott, 2000 | 2>1>0-back, schizophrenia>controls | 7 | 0,1,2 | Numbers |
| Callicott, 2003 | 2-back>0-back, schizophrenics>controls | 7 | 0,2 | Numbers |
| Ettinger, 2011 | 2,1,0 (group by load), schizophrenics>controls | 2 | 0,1,2 | Dots |
| Guerrero-Pedraza, 2012 | 2-back>asterisk, first episode psychosis>controls | 2 | asterisk,2 | Letters |
| Guimond, 2018 | 2-back>0-back, schizophrenia patients>controls | 1 | 0,2 | Faces |
| Habel, 2010 | 2-back>0-back, male schizophrenics>controls | 5 | 0,2 | Letters |
| Jansma, 2003 | 3,2,1>0-back, schizophrenics (atypical)>controls | 7 | 0,1,2,3* | Dots |
| Li, 2019 | 2-back>0-back, schizophrenia patients>controls | 3 | 0,2 | Symbols and Shapes |
| Madre, 2013 | 2>1>asterisk, schizo-affectives>controls | 1 | asterisk,1,2 | Letters |
| Mendrek, 2004 | 2-back>0-back, schizophrenics>controls | 6 | 0,2 | Letters |
| Meyer-Lindenberg, 2001 | 2-back>0-back, schizophrenics>matched controls | 4 | 0,2 | Numbers |
| Nejad, 2011 | 2-back>1-back, first-episode SCZ>controls | 6 | 1,2 | Letters |
| Ortiz-Gil, 2011 | 2-back>1-back, schizophrenics>controls | 2 | 1,2 | Letters |
| Ortiz-Gil, 2011 | 2-back>asterisk, schizophrenics>controls | 2 | asterisk,2 | Letters |
| Pomarol-Clotet, 2008 | 2-back>asterisk, chronic schizophrenia>controls | 2 | asterisk,2 | Letters |
| Pomarol-Clotet, 2010 | 2-back>asterisk, schizophrenic patients>controls | 2 | asterisk,2 | Letters |
| Royer, 2009 | 2-back>0-back, schizophrenia>controls | 4 | 0,2 | Letters |
| Sabri, 2003 | 2-back>0-back, chronic schizophrenia>controls | 9 | 0,2 | Numbers |
| Salgado-Pineda, 2018 | 2-back>1-back, chronic schizophrenia>controls | 18 | 1,2 | Letters |
| Salgado-Pineda, 2018 | 2-back>1-back, FEP schizophrenia>controls | 6 | 1,2 | Letters |
| Sapara, 2013 | 2-back>rest, schizophrenia>controls | 5 | rest,2 | Dots |
| Sapara, 2013 | 2-back>1-back, schizophrenia>controls | 3 | 1,2 | Dots |
| Scheuerecker, 2008 | 2-back>0-back, schizophrenic inpatients>controls | 1 | 0,2 | Letters |
| Scheuerecker, 2008 | 2-back (d)>0-back (d), schizophrenia>controls | 1 | 0d, 2d | Letters |
| Schneider, 2007 | 2-Back>0-back, schizophrenia>controls | 3 | 0,2 | Letters |
| Tan, 2006 | 2-back>1-back, schizophrenia>controls | 2 | 1,2 | Numbers |
| Thermenos, 2016 | 2-back>0-back, CHR for schizophrenia>controls | 2 | 0,2 | Letters |
| Walter, 2003 | 2-back>0-back, schizophrenia>controls | 1 | 0,2 | Letters |
| Yoo, 2005 | 2-back>0-back, schizophrenia>controls | 2 | 0,2* | Faces |
| Schizophrenia < Controls |  |  |  |  |
| Callicott, 2000 | 2>1>0-back, controls>schizophrenia | 7 | 0,1,2 | Numbers |
| Callicott, 2003 | 2-back>0-back, controls>schizophrenics | 7 | 0,2 | Numbers |
| Guimond, 2018 | 2-back>0-back, controls>schizophrenia patients | 1 | 0,2 | Faces |
| Habel, 2010 | 2-back>0-back, controls>male schizophrenics | 6 | 0,2 | Letters |
| Honey, 2003 | 2-back>0-back, controls>schizophrenic patients | 2 | 0,2 | Letters |
| Karch, 2009 | 2-back>0-back, controls>schizophrenic patients | 13 | 0,2 | Numbers |
| Kumari, 2006 | 2-back>0-back, controls>schizophrenia | 1 | 0,2 | Dots |
| Loeb, 2018 | 2-back>1-back, controls>COS | 12 | 1,2 | Letters |
| Loeb, 2018 | 2-back>1-back, siblings>COS | 11 | 1,2 | Letters |
| Madre, 2013 | 2>1>asterisk, controls> schizo-affectives | 3 | asterisk,1,2 | Letters |
| Meisenzahl, 2006 | 2-back>0-back controls>schizophrenic inpatients | 8 | 0,2 | Letters |
| Mendrek, 2004 | 2-back>0-back, controls>schizophrenia | 2 | 0,2 | Letters |
| Meyer-Lindenberg, 2001 | 2-back>0-back, matched controls>schizophrenics | 5 | 0,2 | Numbers |
| Moser, 2018 | 2-back>0-back, schizophrenia>controls | 6 | 0,2 | Objects |
| Ortiz-Gil, 2011 | 2-back>asterisk, controls>schizophrenics | 1 | asterisk,2 | Letters |
| Perlstein, 2001 | GroupXload (0,1,2-back), controls>schizophrenia | 1 | 0,1,2 | Letters |
| Perlstein, 2003 | GroupXload(0,1,2-back), controls>schizophrenia | 1 | 0,1,2 | Letters |
| Pomarol-Clotet, 2008 | 2-back>asterisk, controls>chronic schizophrenia | 3 | asterisk,2 | Letters |
| Pomarol-Clotet, 2010 | 2-back>asterisk, controls>schizophrenic patients | 3 | asterisk,2 | Letters |
| Salgado-Pineda, 2018 | 2-back>1-back, controls>chronic schizophrenia | 36 | 1,2 | Letters |
| Salgado-Pineda, 2018 | 2-back>1-back controls>FEP schizophrenia | 12 | 1,2 | Letters |
| Sapara, 2013 | 2-back>0-back, schizophrenia>controls | 3 | 0,2 | Dots |
| Scheuerecker, 2008 | 2-back>0-back, controls>schizophrenic inpatients | 3 | 0,2 | Letters |
| Scheuerecker, 2008 | 2-back (d)>0-back (d), controls>schizophrenia | 1 | 0d,2d | Letters |
| Schneider, 2007 | 2-Back>0-back, controls>schizophrenia | 2 | 0,2 | Letters |
| Wykes, 2002 | 2-back>0-back, controls>schizophrenia | 10 | 0,2 | Letters |
| Yoo, 2005 | 2-back>0-back, controls> schizophrenia | 15 | 0,2* | Faces |

## RESULTS

### N-back working memory network within each group

To identify brain regions in healthy individuals and in those with psychopathology that increase in activation with increasing working memory load during the n-back, we conducted separate within-group meta-analyses. We first collapsed data across all healthy participants that were included in any of the undermentioned contrasts. Healthy participants activated an array of regions typically associated with prior examinations of working memory during the n-back paradigm (22, 23). These regions included bilateral inferior parietal lobule (IPL) extending to precuneus, bilateral insula, bilateral declive of the cerebellum, bilateral mid frontal gyrus (MFG), left precentral gyrus extending into the anterior cingulate, right superior frontal gyrus (SFG), left ventral anterior nucleus of the thalamus extending to the mediodorsal nucleus, and bilateral tuber of the cerebellum (Figure 1, first row; Table 3).

**Figure 1.**
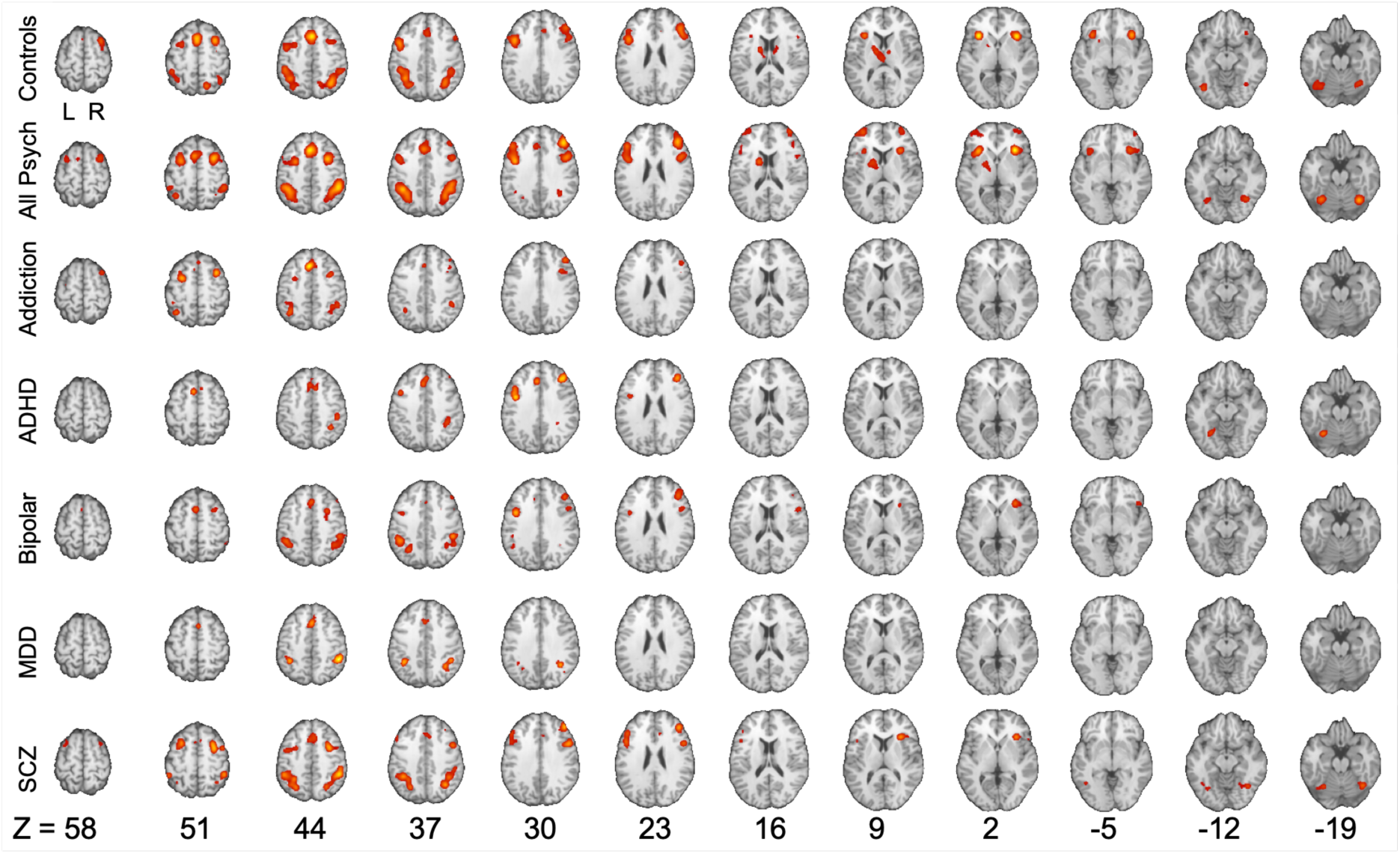
Within-group contrasts. From top to bottom are n-back working memory-related activations in healthy controls, participants with addiction, ADHD, bipolar disorder, major depressive disorder (MDD), and schizophrenia (SCZ). These analyses include data from a total of 160 experiments and 4509 subjects, from which 1603 foci were obtained.

**TABLE 3.** RESULTS

| Condition/Cluster | Volume (mm<sup>3</sup>) | Peak Coordinates (x, y, z) | Label | Brodmann area |
| --- | --- | --- | --- | --- |
| **Addiction** |  |  |  |  |
| Cluster 1 | 6728 | 37.5, 15.7, 40 | Right MFG | 6/9 |
| Cluster 2 | 3672 | -1.4, 21.5, 44.1 | Left Cingulate Gyrus | 8 |
| Cluster 3 | 3648 | -35.2, -53.2, 45.3 | Left IPL | 40/7 |
| Cluster 4 | 2984 | -25.7, 0.5, 50.2 | Left MFG | 6 |
| Cluster 5 | 2848 | 40.5, -46.8, 41.6 | Right IPL | 40 |
| **ADHD** |  |  |  |  |
| Cluster 1 | 5976 | -1.3, 21.1, 41 | Left MFG | 32/6 |
| Cluster 2 | 3896 | -41.2, 5.9, 30.3 | Left precentral gyrus | 9 |
| Cluster 3 | 3504 | 37.6, -41, 39.6 | Right IPL | 40 |
| Cluster 4 | 3296 | 38.6, 32.9, 27.8 | Right MFG | 9 |
| Cluster 5 | 3096 | -26.8, -60.9, -18.7 | Left Declive | * |
| **Bipolar Disorder** |  |  |  |  |
| Cluster 1 | 7000 | -38.8, -48.7, 38.7 | Left IPL | 40 |
| Cluster 2 | 6368 | 45.3, -44.5, 40.9 | Right IPL | 40 |
| Cluster 3 | 3608 | 41.9, 32, 26.4 | Right MFG | 9 |
| Cluster 4 | 3264 | -1.1, 14.7, 46.1 | Left Cingulate Gyrus | 6 |
| Cluster 5 | 3016 | -40, 3.7, 29.9 | Left Precentral Gyrus | 6 |
| Cluster 6 | 2848 | 34.5, 18, 1.1 | Right Insula | 47 |
| Cluster 7 | 2536 | 48, 9.3, 24 | Right IFG | 9 |
| Cluster 8 | 2352 | 27.9, 4.8, 46.5 | Right MFG | 6 |
| **Controls** |  |  |  |  |
| Cluster 1 | 11976 | -41.3, 8.2, 34.1 | Left MFG | 6 |
| Cluster 2 | 11760 | 33.3, -55.8, 42.9 | Right IPL | 7/40 |
| Cluster 3 | 11608 | -36.5, -50.3, 41.4 | Left IPL | 40 |
| Cluster 4 | 8400 | 0.1, 16.6, 45.1 | Right MFG | 6/32 |
| Cluster 5 | 7152 | 45.2, 24.8, 26.3 | Right MFG | 9 |
| Cluster 6 | 6424 | -35, -65, -21 | Left Declive | * |
| Cluster 7 | 5552 | -4.8, -9.8, 11.5 | Left Thalamus (MD) | * |
| Cluster 8 | 4560 | 30.8, 8.7, 51.6 | Right MFG | 6 |
| Cluster 9 | 4448 | -30.9, 19.1, 2.2 | Left Insula | 13 |
| Cluster 10 | 4224 | 32.5, 19.4, -1 | Right Insula | 13/47 |
| Cluster 11 | 3808 | 34.6, -62.6, -23 | Right Declive | * |
| **Depression** |  |  |  |  |
| Cluster 1 | 5496 | 41.1, -48.4, 39.2 | Right IPL | 40 |
| Cluster 2 | 3144 | -35.1, -49.4, 38.2 | Left IPL | 40 |
| Cluster 3 | 2920 | 1.1, 17.6, 43.3 | Right Cingulate Gyrus | 32 |
| **Psychopathologies** |  |  |  |  |
| Cluster 1 | 23424 | 38, 20.6, 32.1 | Right MFG | 6/9 |
| Cluster 2 | 22176 | -36.7, 11.1, 28.9 | Left MFG | 6/9 |
| Cluster 3 | 14032 | 40.9, -47.1, 41.1 | Right IPL | 40 |
| Cluster 4 | 13144 | -35.8, -50.4, 40.9 | Left IPL | 40 |
| Cluster 5 | 10528 | -0.3, 18.8, 43.4 | Left MFG/Cingulate | 6/32 |
| Cluster 6 | 5304 | 32.6, 20.1, 1.4 | Right Insula | 13 |
| Cluster 7 | 4648 | 33.4, -62.7, -19.3 | Right Cerebellar Declive | N/A |
| Cluster 8 | 4504 | -15.2, -2.8, 9.7 | Left Lentiform Nucleus | N/A |
| Cluster 9 | 4456 | -30.5, -62.8, -21.2 | Left Cerebellar Declive | N/A |
| Cluster 10 | 4392 | -34.3, 49.6, 7.8 | Left MFG | 10 |
| Schizophrenia |  |  |  |  |
| Cluster 1 | 9448 | 40.4, -47.7, 42.5 | Right IPL | 40 |
| Cluster 2 | 9424 | -35.1, -51.2, 42 | Left IPL | 40 |
| Cluster 3 | 8992 | 37.8, 5.9, 41.3 | Right MFG | 6 |
| Cluster 4 | 5320 | -45.8, 18.6, 25.3 | Left MFG | 9 |
| Cluster 5 | 4344 | -29.8, 5.5, 48.9 | Left MFG | 6 |
| Cluster 6 | 3904 | 3.3, 19.6, 41.4 | Right Cingulate Gyrus | 32 |
| Cluster 7 | 2952 | 40.7, 36.3, 26.7 | Right MFG | 9 |
| Cluster 8 | 2936 | 34.4, 20.6, 6.2 | Right Insula | 13 |
| Cluster 9 | 2784 | -32.8, -61.7, -17 | Left Declive | * |
| Cluster 10 | 2744 | 35.7, -60.7, -17.2 | Right Declive | * |
| Controls > Addiction |  |  |  |  |
| Cluster 1 | 2096 | -35.3, -64.7, 37.2 | Left Precuneus | 39 |
| Controls > ADHD |  |  |  |  |
| Cluster 1 | 2416 | 39.6, 13.3, 34.8 | Right MFG | 9 |
| Controls > Bipolar |  |  |  |  |
| Cluster 1 | 2104 | -32.5, -0.2, 48.3 | Left MFG | 6 |
| Controls > Depression |  |  |  |  |
| None |  |  |  |  |
| Controls > Schizophrenia |  |  |  |  |
| Cluster 1 | 2576 | 4.1, - 58.1, -16.9 | Right Culmen | * |
| Cluster 2 | 2224 | 32.5, 23.4, -2.9 | Right IFG | 47 |
| Addiction > Controls |  |  |  |  |
| None |  |  |  |  |
| ADHD > Controls |  |  |  |  |
| None |  |  |  |  |
| Bipolar > Controls |  |  |  |  |
| Cluster 1 | 4112 | -1.2, 38.3, -13.7 | Left MFG | 11 |
| Depression > Controls |  |  |  |  |
| None |  |  |  |  |
| Schizophrenia > Controls |  |  |  |  |
| Cluster 1 | 6000 | -2, 32.3, -8.2 | Left anterior cingulate | 11/32 |
| Psychopathologies > Controls |  |  |  |  |
| Cluster 1 | 11720 | -2, 34.2, -7.7 | Left anterior cingulate | 11/32 |
| Controls > Psychopathologies |  |  |  |  |
| Cluster 1 | 4480 | 5.2, -61, 46.5 | Right Precuneus | 7 |
| Cluster 2 | 2928 | 33.5, 20.5, -2.8 | Right Insula | 13 |

We furthermore collapsed data across all within-group psychopathologies contrasts. We found convergence of activation in a breadth of brain regions, including bilateral SFG, bilateral MFG, bilateral IPL, right IFG, bilateral cerebellar declive, left precentral gyrus, left claustrum, left cingulate gyrus, right insula, left caudate, and left putamen (Figure 1, second row; Table 3).

Across many of the psychopathologies the topology of the ALE maps related to an increase in activation with increasing working memory load was consistent with that observed in the healthy participants and each other. For example, when considering individuals with addiction, increased working memory load was associated with increased activation in left cingulate gyrus, left sub-gyral (Brodmann area [BA] 6), bilateral superior parietal lobule (SPL), bilateral IPL, bilateral MFG, and bilateral precentral gyrus (Figure 1, third row; Table 3).

In the ADHD group, the n-back working memory network was characterized by activation in left SFG, left cingulate gyrus, right medial frontal gyrus (MeFG), left precentral gyrus, bilateral MFG, right IPL, right sub-gyral (BA 40), and left cerebellar declive (Figure 1, fourth row; Table 3).

In individuals with Bipolar Disorder, increased working memory load resulted in increased activation in bilateral IPL, left angular gyrus, bilateral MFG, left MeFG, left SFG, left cingulate gyrus, bilateral precentral gyrus, right claustrum, and right inferior frontal gyrus (IFG) (Figure 1, fifth row; Table 3).

In MDD, increased working memory load was associated with increased activation in bilateral IPL, right superior temporal gyrus (STG), bilateral precuneus, left angular gyrus, and left MFG (Figure 1, sixth row; Table 3).

In the schizophrenia group, increased working memory load during the n-back task was associated with activation in bilateral IPL, right angular gyrus, bilateral SPL, bilateral MFG, bilateral IFG, bilateral precentral gyrus, right cingulate gyrus, left MeFG, right anterior cingulate, right SFG, right insula, left inferior temporal gyrus (ITG), left fusiform gyrus, bilateral cerebellar declive, and left cerebellar culmen. (Figure 1, sixth row; Table 3).

### Greater working memory-related activations in healthy individuals

To evaluate what brain regions showed greater working memory-related activations in healthy individuals compared to psychopathology more generally and maximize our power to detect an effect, we collapsed data across all between-group contrasts (Healthy Controls > Psychopathology) for the subsequent meta-analysis. Healthy individuals demonstrated greater activation in bilateral precuneus and right IFG (triangularis/opercularis) extending into insula (Figure 2, Table 3). Healthy participants from all studies contributed to the formation of the bilateral precuneus cluster, while subjects from MDD and schizophrenia studies contributed to the formation of the right IFG cluster.

**Figure 2.**
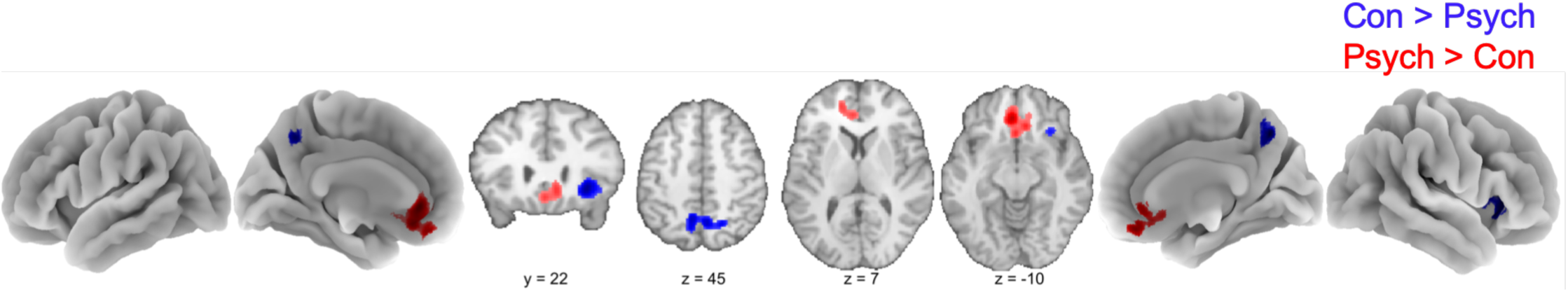
Between-group contrasts. Clusters show regions in which healthy controls exhibit greater activation than participants with psychopathologies (blue) or regions in which participants with psychopathologies demonstrate greater activation that healthy controls (red). The latter contrast converges in lACC/mPFC extending into OFC, while the prior contrast exhibits two clusters of convergence in precuneus and rIFG extending into insula. Darker colors indicate higher ALE values (and thus lower *p*-values). Con = Controls. Psych = All psychopathologies.

Follow-up analyses examined the same comparisons – brain regions that exhibit greater n-back working memory-related activation in healthy subjects compared to those with a psychiatric diagnosis – but for each psychopathology separately. Across addiction studies, healthy subjects exhibited greater working memory-related activation than patients in left precuneus. Compared to individuals with ADHD, healthy controls exhibited greater activation in right MFG. Relative to those with bipolar disorder, healthy controls exhibited greater activation in left MFG. No brain regions exhibited significantly greater activation in healthy controls compared to individuals with MDD. Right IFG, insula, cerebellar culmen, nodule, and declive all exhibited greater activation in healthy controls compared to individuals with schizophrenia on the n-back task (Figure 3, Table 3).

**Figure 3.**
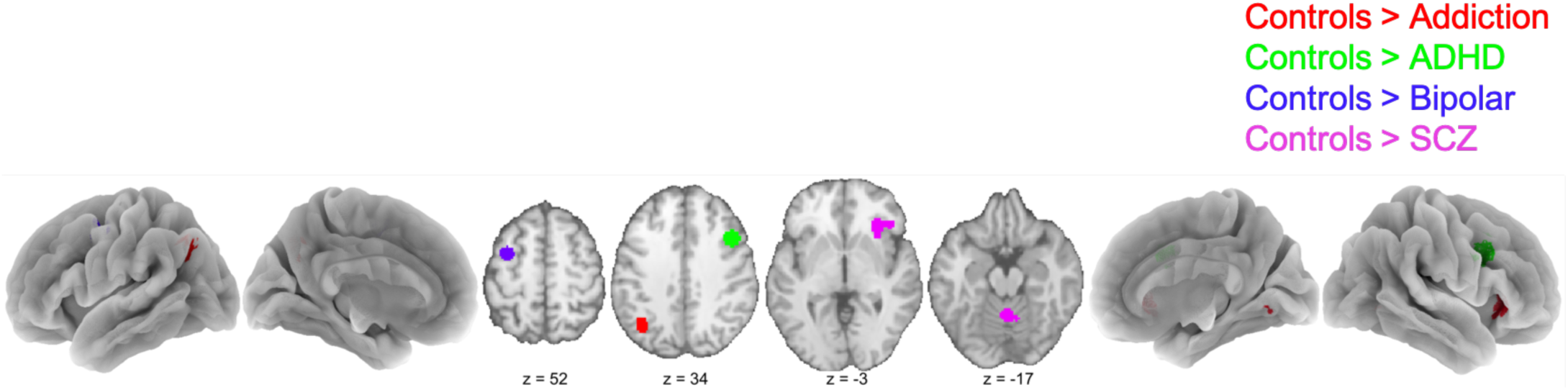
Between-group contrasts. Each contrast represents regions of n-back-related hyperactivation in healthy controls relative to participants with addiction (red), ADHD (green), bipolar disorder (blue), or schizophrenia (SCZ; pink). These regions included left precuneus, right MFG, left MFG, as well as right IFG (extending into insula) and cerebellum, respectively. No significant clusters were obtained for participants with major depressive disorder.

### Greater working memory related activations in individuals with psychopathology

We conducted an additional meta-analysis to examine effects in which patients across psychopathologies exhibited greater activations relative to controls during the n-back task (i.e. Psychopathology > Healthy Controls). We observed a significant convergence of hyperactivation across all psychopathologies compared to controls in the left anterior cingulate cortex/medial prefrontal cortex (lACC/mPFC) (Figure 2, Table 3). This cluster (-4, 36, -2) was centered in a central or hub region of the default mode network (DMN), a group of brain regions that are more active during rest than during task performance (24, 25). Experiments from the addiction, ADHD, bipolar disorder, MDD, and schizophrenia greater than controls contrasts contributed to the formation of this cluster.

We next conducted a follow-up meta-analysis to examine effects in which patients across individual psychopathologies exhibited greater activations relative to controls during the n-back task. The ALE algorithm yielded no significant regions of convergence where participants diagnosed with addiction, ADHD, or MDD exhibited greater activation during the n-back working memory task compared to controls. In contrast, patients diagnosed with bipolar disorder and schizophrenia exhibited greater activation in the lACC extending into the mPFC during the n-back task compared to healthy controls (Figure 4, Table 3).

**Figure 4.**
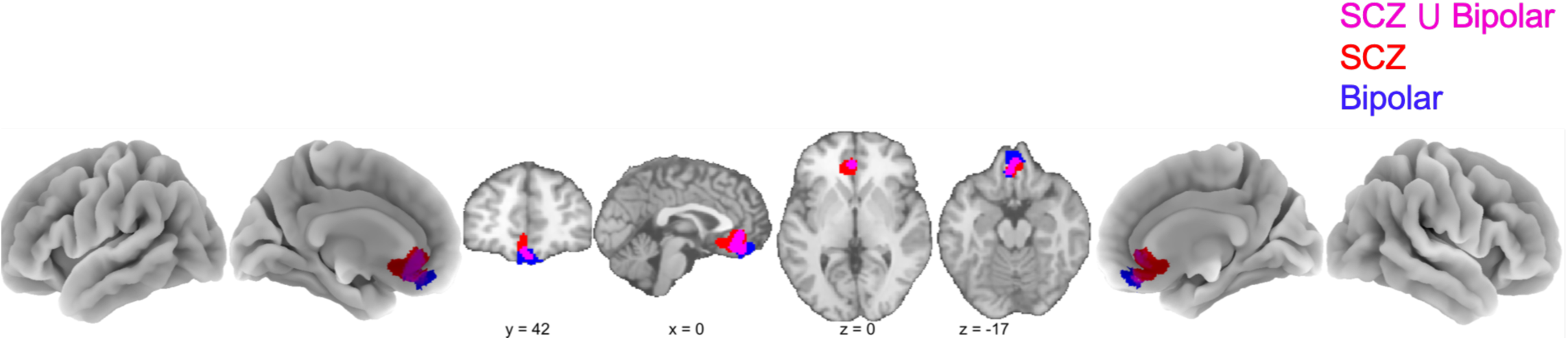
Between-group contrasts. Contrasts represent a region in the brain in which hyperactivity was observed in participants with schizophrenia (SCZ; red) or bipolar disorder (blue). These regions converge (pink) in lACC/mPFC, with a slight extension into orbitofrontal cortex.

## DISCUSSION

Deficits in working memory are commonly reported in individuals with a variety of psychopathologies. We examined neuroimaging data across ADHD, addiction, bipolar disorder, MDD, and schizophrenia to identify similarities and differences in human brain activation during the same working memory paradigm (n-back task). Elucidating convergent and/or divergent neurobiological correlates may shed light on their underlying pathophysiology (1–6). With increasing working memory load participants with psychiatric disorders when compared to controls exhibited hyperactivity in the mPFC, a hub of the DMN, while controls showed greater activation in the right IFG and bilateral precuneus. These results provide novel and compelling evidence that in addition to frontal and parietal dysfunction, DMN intrusion may constitute a conserved mechanism of dysregulation across psychopathologies resulting in poorer working memory performance.

Patterns of activation with increasing working memory load in each group were remarkably consistent following our within-group meta-analyses. Prototypical regions – including the bilateral IPL, bilateral MFG, bilateral anterior insula, and SFG – extending ventrally into the anterior cingulate cortex – were seen within each group and are consistent with prior meta-analytic studies examining working memory load in non-patient populations (22, 23). The lack of grossly discernable differences is not entirely unexpected given that impairments in behavior often result in subtle, rather than large scale, changes in functional activity. While some qualitative differences in the topology of ALE activations were evident across groups, we believe these dissimilarities arise from differences in power across contrasts. For example, the within-group psychopathologies contrast had the most statistical power (106 experiments, 1042 foci); thus, it follows that the working memory network for this contrast should be more robust than in the MDD contrast, which had the least power (15 experiments, 142 foci). Finally, we note that while there are observable differences in the within-group, collapsed controls contrast versus psychopathologies contrast (for example, in bilateral MFG), regions of spatial convergence or divergence do not represent direct statistical comparisons. To accomplish this, meta-analyses should be conducted on publications with between-group statistical comparisons that include both patient and healthy control samples. Accordingly, meta-analyses comparing between-group differences are essential for identifying convergence and divergence of activations related to working memory impairments in psychiatric disorders.

To directly examine functional differences between healthy controls and subjects with psychopathology as working memory load increases, we performed meta-analyses of between-groups data. These analyses detailed regions in healthy subjects that exhibited greater working memory load-related activation relative to those with psychopathology, and vice versa. Considerable heterogeneity in activations were evident across groups which likely reflects differences in power across the different psychopathologies (as noted above). However, it is possible that the observed differences may highlight unique functional impairments for each psychopathology when compared to healthy individuals. We focus our discussion, rather, on the undermentioned collapsed data, which sheds insight into common pathophysiology across all disease states and is bolstered by greater statistical power.

To examine common neurobiological correlates, we first collapsed across all contrasts to examine brain regions in which healthy subjects exhibited greater working memory load-related activation than individuals with any given psychopathology. With this analysis, we were able to examine brain regions that exhibited hyperactivity in healthy controls relative to individuals with mental illness, who are typically impaired on the n-back task. We found convergence of activation in bilateral precuneus and right IFG extending into insula. The precuneus has long been implicated in successful episodic memory retrieval (26, 27), and is associated with better performance on spatial working memory tasks (28, 29). Further, it has been shown to activate during the n-back in healthy subjects regardless of memory load, object, age, or gender (23). Differential recruitment of bilateral precuneus in this instance requires further investigation, though we posit that its greater recruitment is coincident with normal cognitive processing during this task. The IFG/insula region is a component of the salience network, a group of brain regions responsible for orienting toward behaviorally salient external events (30, 31). While insula is implicated in disparate cognitive responses such as emotional and interoceptive processing, in this instance it is likely responsible for orienting toward salient stimuli and switching between networks (DMN and central executive) to permit access to attention and working memory stores (32, 33). We thus assert that, in healthy participants, greater insula recruitment likely reflects an “open door policy” for working memory that is ‘shut’ in those with psychopathology.

While working memory impairments may arise from aberrant salience detection, behavioral performance may also suffer due to spurious activations of regions unrelated to the task at hand. Supporting this possibility, individuals with psychopathology compared to healthy controls during the n-back task exhibited greater activation with increasing working memory load in the mPFC. When data from all psychopathologies was collapsed, the cluster was comprised of approximately 14.8% MDD studies, 37.0% Bipolar studies, 33.3% schizophrenia studies, 11.11% addiction studies, and 3.7% ADHD studies. When examining each psychopathology separately subjects with schizophrenia and bipolar disorder both exhibited a similar cluster that survived corrections for multiple comparisons, while data from ADHD, addiction, and MDD groups did not survive statistical thresholding. Thus, pooling across psychopathologies helped identify the ADHD, addiction, and MDD groups as contributors to the mPFC cluster, while on their own not significant.

The mPFC is critical for a diverse array of functions in the brain. Lesions in this region are associated with drastic impairments in personality, affect, emotion, decision-making, and general cognition (34). Literature linking aberrant mPFC function to psychiatric disorders is replete. For example, structural analyses suggest that mPFC-amygdala white matter connectivity predicts anxiety and depressive symptoms in childhood (35), and functional connectivity between these regions is negatively correlated with PTSD symptoms (36). Smaller mPFC volume in adolescents predicts ADHD symptoms after 5 years, while mPFC activity in individuals with schizophrenia and comorbid nicotine addiction (relative to healthy controls) is enhanced following exposure to cigarette cues (37). Finally, a recent meta-analysis supports the assertion that distinct subregions of the mPFC are associated with psychopathologies such as PTSD, addiction, depression, social anxiety, and schizophrenia (38). Of note, the mPFC plays an integral role in memory; indeed, those with mPFC lesions are prone to memory confabulations, poor schematic memory, and impaired environmental context effects on memory formation (34). mPFC-hippocampal interactions have been shown to mediate memory-based decision-making (39, 40) as well. Further, psychophysiological analyses during a working memory task support the role of mPFC as an “emotional gating” mechanism in instances of high cognitive load (41). Indeed, this supports the earlier thesis that mPFC connectivity facilitates emotion-cognition interactions, or simply, the interplay between affect and reason (42).

In addition to being a key region responsible for the integration of cognitive and emotional stimuli, the mPFC is also a known hub of the DMN – an organized network of brain regions that are more active during rest than during cognitive tasks (24, 43). Intrusion of the DMN during cognitive tasks may reflect insufficient top-down attentional control, leading to performance decrements (44). This “default mode interference hypothesis” suggests that spontaneous low frequency activity in regions of the DMN, such as the mPFC, can emerge during the performance of a task and occupy neural resources necessary for performing that task, ultimately resulting in behavioral impairments (45). Strikingly, Whitfield-Gabrieli et al. (46) examined patients with schizophrenia and first-degree relatives of those with schizophrenia, and found that those particular individuals exhibited reduced task-related suppression of a similar, though more anterior, region in mPFC during an n-back working memory paradigm. Whitfield-Gabrieli et al. (46) was not incorporated in our analyses as the data did not conform to our inclusion criteria. Our result provides further replication for the role of dysregulated mPFC activity as a direct contributor to poor working memory performance in schizophrenia and bipolar disorder, and suggests that it may have a role to play in ADHD, addiction, and MDD as well.

### Conclusion

We have found evidence for greater recruitment of regions within the salience network (i.e. IFG and insula) in healthy individuals along with DMN (i.e. mPFC) intrusion in psychiatric patients during performance of the n-back working memory paradigm. Not only do these results provide evidence for the default mode interference hypothesis, they also speak more generally in support of the triple network model of Menon (47) which posits that aberrant function within three neurocognitive networks constitute a common feature among multiple psychopathologies. These three networks, the frontoparietal central executive network (CEN), salience network (SN), and default mode network (DMN) have been shown to be dysregulated in schizophrenia, depression, dementia, autism, and anxiety (47). We extend these findings to suggest that disruption in a combination of at least two of these networks, the DMN and SN, play a role in affecting working memory performance of individuals with schizophrenia and bipolar disorder, and that such a role may exist in ADHD, addiction, and MDD as well.

### Limitations and Future Directions

In meta-analyses the recommended number of experiments per contrast is 20 (Eickhoff, personal communications). Here, we took great efforts to meet this recommendation, while at the same time stringently applying our inclusion/exclusion criteria to obtain the most accurate and interpretable results. On average, we had 30.21 experiments per contrast including pooled contrasts such as ‘All Psychopathologies’ ‘Controls’, and ‘Psych > Con’. With pooled contrasts excluded, there were 17.33 experiments per contrast. With this in mind, however, the minimum number of experiments included was 8 in the Con > Addiction contrast. Caution should be taken in over-interpreting the related results. In fact, between-group contrasts of each psychopathology separately, which are the most informative to pathologically related differences in brain activation, had the fewest number of experiments on average (mean = 15.4). This notable limitation speaks to the need for replication studies and large sample sizes to facilitate the examination of between group differences. However, our approach to collapsing across psychopathologies was sufficiently powered and provided mechanistic insight to working memory impairments. As this meta-analysis captures cross-sectional data, further studies are necessary to determine whether the observed differences in brain activity are a cause or consequence of subjects’ respective diagnoses. Finally, recent research has used the mPFC as a target for fMRI-based neurofeedback (48). Future studies should determine whether this approach is capable of mitigating working memory deficits in individuals with psychopathology.

## ACKNOWLEDGEMENTS

We would like to thank Drs. Simon Eickhoff and Mick Fox for their help in implementing the ALE algorithm, as well as Dr. Michael Cody Riedel for help with visualization. We would finally like to thank Candelaria Albarracin for help with data collection. This work was supported by startup funds provided by the Florida International University to A.T.M and by NIH R01-DA041353 (A.R.L.).

## AUTHOR CONTRIBUTIONS

Conceptualization, A.T.M. and M.C.F.; Investigation, M.C.F.; Formal Analysis, M.C.F.; Writing – Original Draft Preparation, M.C.F.; Writing – Review and Editing, M.C.F., A.R.L., and A.T.M.; Supervision, A.T.M.; Funding Acquisition, A.T.M and A.R.L.

